# Integrative chromatin state annotation of 234 human ENCODE4 cell types using Segway reveals disease drivers

**DOI:** 10.1101/2023.10.26.564254

**Authors:** Marjan Farahbod, Abdul Rahman Diab, Paul Sud, Meenakshi Kagda, Ian Whaling, Mehdi Foroozandeh, Ishan Goel, Habib Daneshpajouh, Benjamin Hitz, J. Michael Cherry, Maxwell Libbrecht

## Abstract

Towards the goal of identifying functional elements in the human genome, the fourth and final phase of the ENCODE consortium has newly profiled hundreds of human tissues using sequencing-based measurements of genomic activity such as ChIP-seq measures of transcription factor binding and histone modification. Chromatin state annotations created by segmentation and genome annotation (SAGA) methods such as Segway have emerged as the predominant integrative summary of such epigenomic data sets. Here, we present the ENCODE4 catalog of Segway annotations, a set of sample-specific genome-wide Segway chromatin state annotations for 234 ENCODE human biosamples inferred from 1,794 functional genomics experiments. We define an updated vocabulary of chromatin state terms that includes patterns of activity present only in a subset of samples or identified only with rarely-performed assays. We show that these ENCODE4 Segway annotations accurately capture both general and cell-type-specific regulatory patterns, and do so with substantially improved sensitivity relative to prior large-scale chromatin annotation sets. This catalog facilitates the downstream discovery of regulatory mechanisms which underlie diseases and traits identified by genome-wide association studies.

## Introduction

Identifying functional elements in the genome is critical to understanding human biology and disease. To that end, the ENCODE consortium has engaged in large-scale mapping of human epigenomes using sequencing-based measurements of genomic activity, including ChIP-seq measurement of transcription factor binding and histone modifications and DNase-seq and ATAC-seq measurement of chromatin accessibility ^1–3^. The fourth and final phase of ENCODE has greatly expanded the set of profiled biosamples. Epigenome mapping facilitates the creation of reference genome annotations, which enable researchers to understand genomic activity in any of the hundreds of characterized cell and tissue types and thus understand the influence of genomic activity on disease and other phenomena ^4,5^.

The ENCODE Encyclopedia is a collection of reference annotations that encompass all outputs of the consortium, with the aim that these annotations provide a resource to the research community. Here, we present a component of this Encyclopedia, the ENCODE4 Segway catalog of chromatin state annotations. Chromatin state annotations are the predominant form of integrated genome annotation ^6^. They are created by segmentation and genome annotation (SAGA) methods such as Segway, ChromHMM and IDEAS ^6–12^ (reviewed in Libbrecht et al. 2021 ^6^). These methods take as input a collection of epigenomic data sets from a given biosample, which may be a primary tissue sample or cell line, and produce an annotation of chromatin states in the genome. Each chromatin state corresponds to a pattern of epigenomic activity such as patterns of transcription, Polycomb repression, or patterns associated with promoters, enhancers or other types of genomic regulatory elements. SAGA methods are unsupervised in that they identify patterns of epigenomic activity through learning a probabilistic model without predefined categories of genomic elements. A researcher must interpret these learned patterns to map them to known categories of genomic functions ^7^.

The ENCODE4 Segway catalog comprises chromatin state annotations for 234 samples created through a SAGA pipeline based around the Segway model ^13^. This pipeline has two main advantages relative to previous approaches (reviewed in Supplementary Material) for performing chromatin state annotation. First, the model incorporates genomic signal strength, as measured by normalized read count, avoiding a binarizing step. Thus Segway chromatin state annotations can distinguish high-signal from low-signal elements. As we show below, doing so greatly improves the model’s sensitivity, allowing it to identify a large number of regulatory elements and in turn increasing its ability to capture gene regulation.

Second, this pipeline involves training an independent Segway model for each sample. This independent approach, in contrast to the alternative “concatenated” approach ^5^ of using a single model for all samples, allows the pipeline to incorporate all data sets available for each sample. To handle the added challenge of interpreting independent models, we used an automated interpretation process as described. Doing so also obviates the need to resort to imputing unperformed assays ^14–17^. While imputation can reduce noise, it creates the risk that annotations may be biased by data observed in other samples ^7,15^. In particular, imputation tends to drive all samples toward a single average and thus can make it harder to identify sample-specific activity ^4,9^. The sample-specific SAGA model can identify epigenetic patterns specific to each sample, and is not at risk of modeling artifacts introduced by applying the sample model to datasets exhibiting experiment-specific patterns. To allow this independent modeling approach to scale, we employ an automated interpretation process which assigns a controlled vocabulary of common chromatin state descriptor terms (e.g. “Promoter”, “Enhancer”) to each identified Segway state.

The resulting reference chromatin state annotations encompass 234 samples and are inferred from 1,794 functional genomics experiments. We define an updated vocabulary of chromatin state terms that includes patterns of activity present only in a subset of samples or identified only with rarely-performed assays. We show that these annotations accurately capture genome biology, and that they do so with enhanced accuracy relative to existing reference chromatin state annotations. We demonstrate they can be used to accurately identify causal genomic elements and cell types underlying associated genetic variation.

## Results

### Genome annotation for 234 samples using Segway

We obtained Segway annotations for each of the samples independently, using a two-step annotation pipeline (Methods, Figure 1A,B,F). In the first step, we applied Segway, which partitions the genome based on data from epigenomic tracks and assigns a state label to each segment such that sections with the same label share similar patterns in the epigenomic tracks (Methods). At this stage, the state labels assigned by Segway solely distinguish different regions and do not bear any biological or functional meaning. We determined the number of labels for each sample according to the count of input tracks, following the previous work ^7^ (Methods); each of the 234 samples has 6-12 input tracks and 14-16 labels.

**Figure 1:**
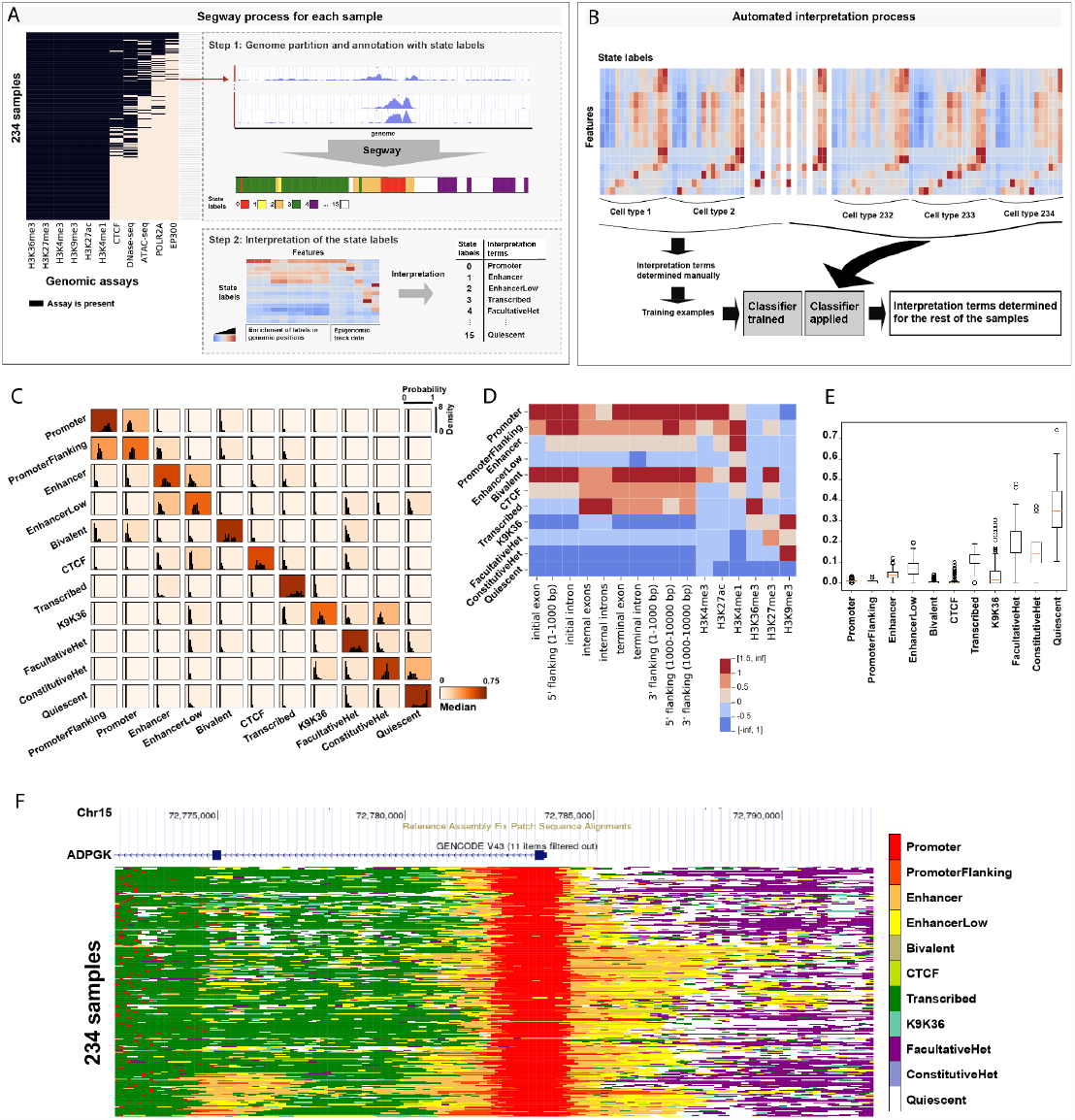
(A) Top left: matrix of input data sets. For each sample, we trained an independent Segway model, then used that model to annotate the genome of that cell type (Methods). We used an automated process to assign a controlled vocabulary of chromatin state descriptor terms to each Segway state (Methods). (B) The automated interpretation process defines a vector of features defining the properties of a given state that researchers usually use for manual interpretation, such as the association with the input data sets and enrichment around annotated genes. We manually interpreted a subset of 90 states to form a training set, then trained a multi-label random forest to recapitulate this interpretation process for the other 3,408 states (Methods). (C) Accuracy of the automated interpreter. Each box shows the distribution of the probabilities for each of the interpretation terms; background color denotes median probability. The term with the highest probability is the output of the classifier for each of the state labels. (D) Association between interpretation terms (vertical) and features input to the interpretation model (horizontal). Color indicates z-score. (E) Distribution of genomic coverage for each interpretation term. Boxplot shows the distribution for a given interpretation term across the 234 samples. (F) Annotations of an example locus in chromosome 15. Vertical axis indicates sample and color indicates annotation label.

In the second step, we map the state labels to the interpretation terms based on the features from each of the state labels. As Segway is an unsupervised model, its identified state labels must be mapped to human-recognizable interpretation terms (e.g. ‘Enhancer’; these are also known as “mnemonics”). In the past, this interpretable process was performed manually by evaluating each label’s pattern to input tracks and relationship to known genomic elements.

Here, we used a recently-developed automated interpretation process to obtain an unbiased and fast mapping of the interpretation terms to biological state labels ^7^. To do so, we manually interpreted a subset of labels and used this subset as a training set to train a random forest classifier to interpret the remaining labels (Figure 1B, Methods). The classifier takes as input a set of 16 features for each Segway state label and outputs one of 11 interpretation terms (Figure 1B, Table 1). Six of the features encode the mean signal of six different histone modification marks in label-associated regions. The remaining 10 features quantify the enrichment of the label at various regions relative to gene bodies (Figure 1D, Methods).

**Table 1:** Vocabulary of interpretation terms for ENCODE4 Segway states.

|  |  |
| --- | --- |
| **Promoter** | Presence of promoter-associated marks H3K4me3 and H3K9ac. Highly enriched at transcription start sites (TSSs). |
| **PromoterFlanking** | Presence of promoter-associated marks H3K4me3 and H3K9ac, but at lower levels than Promoters. Tends to occur upstream or downstream of TSSs. |
| **Enhancer** | Presence of the enhancer-associated marks H3K27ac and H3K4me1. |
| **EnhancerLow** | Same as Enhancer, but with lower signal values. |
| **Bivalent** | Presence of both activating (H3K27ac) and repressive (H3K27me3) marks. |
| **CTCF** | Presence of the transcription factor CTCF, thought to play a role in chromatin conformation. In samples without measured CTCF, this label marks positions with CTCF-associated marks. |
| **Transcribed** | Characterized by the transcription-associated mark H3K36me3. Highly enriched in annotated gene bodies. |
| **K9K36** | Presence of the marks H3K9me3 and H3K36me3, a pattern associated with zinc finger genes. |
| **FacultativeHet** | Facultative (Polycomb) Heterochromatin, characterized by the histone modification H3K27me3. Thought to carry out cell type-specific repression. |
| **ConstitutiveHet** | Constitutive Heterochromatin, characterized by the histone modification H3K9me3. Marks permanently silent regions such as centromeres and telomeres. |
| **Quiescent** | Lack of any marks |

We developed a new vocabulary of 11 interpretation terms in order to capture the diversity of epigenomic patterns present in the 234 samples (Table 1). This vocabulary identifies regulatory regions using six terms: Promoter, PromoterFlanking, Enhancer, EnhancerLow, Bivalent and CTCF; inactive heterochromatin using three terms: FacultativeHet, ConstitutiveHet and Quiescent, as well as a term for transcribed genes (Transcribed) and one for an uncommon combination of histone marks associated with zinc finger genes (K9K36). We should highlight that each term describes an empirically-observed pattern of epigenetic activity and not a functional hypothesis. For example, “Enhancer” labels mark segments with enhancer-associated epigenetic marks but we do not intend to assert that all such segments act as functional enhancers ^6^. Similarly, the distinction between Enhancer and EnhancerLow describes high versus low epigenetic read counts, not necessarily high versus low strength of regulatory function.

We found that the interpreter accurately recapitulates manual labeling (Figure 1C, SFigure1). When the interpreter disagrees with manual interpretation, the disagreement occurs between similar terms (Figure 1C, SFigure 3). For example, Promoter and PromoterFlanking are often switched, but rarely e.g. Promoter and Quiescent. However, we also found that a confused interpreter is a good indicator of poor data quality for a given sample (SFigure04).

Specifically, samples with low posterior probabilities of label assignment from the interpreter tend to be dominated by inactive labels in other annotations based on the same data, suggesting poor data quality (Pearson correlation -0.37 between median classifier posterior and inactive coverage). Thus interpreter posterior can be used as a quality control metric.

Based on this, we identified a list of 16 low quality samples (Supplementary Table ST04). Although our interpretation process does not explicitly require that each sample include labels with all the different interpretation terms, most samples do so. Automatic interpretation of Segway annotations identifies all the labels active promoter, enhancer, and transcribed regions and inactive FacultativeHet, ConstitutiveHet, and Quiescent regions in almost all of the samples (> 94%). Only a subset of samples include Bivalent and K9K36 labels (SFigures 5,6), likely because these chromatin states are not present in all cell types. Furthermore, CTCF ChIP-seq data is present in only 51% of samples and thus the corresponding label is present in only a subset (32%) of samples. Also note that the interpreter does not take CTCF signal as input, yet it is still able to reliably identify CTCF labels based on its associated features (SFigure 1).

Our annotations capture a large amount of previously unannotated activity. Our annotations label 12-48% (median 27%) of the genome with one of the 8 active labels (Figure 1E, SFigure 2), mostly due to Transcribed and EnhancerLow labels. For comparison, the largest existing set of reference chromatin state annotations is the EpiMap reference generated in Boix et al. 2021 ^20^; these annotations assign 35-92% (median 72%) of the genome to the uninformative Quiescent label (SFigure 2). In contrast, our annotations assign just 10-75% (median 35%) as Quiescent. Instead, they label on average an additional 12% and 15% as the more-specific heterochromatin types ConstitutiveHet and FacultativeHet, and label on average an additional 9% as one of the 8 active types. This large increase in sensitivity is likely due to Segway’s use of genomic signals as opposed to binarized peak calls.

## Annotations accurately capture regulatory activity and transcription

We found that Segway annotations accurately distinguish expressed versus silent genes (Figure 2A). Highly-expressed genes tend to have the Promoter label at their transcription start site (TSS) and the Transcribed label throughout their gene bodies. Conversely, silent genes more often have the Bivalent label at their promoter and their gene bodies are enriched for FacultativeHet and show no particular enrichment for the Transcribed state. Genes expressed at a low level are disproportionately labeled with Bivalent and are only slightly enriched for the Transcribed label. Our annotations show a clear difference between the enrichment of labels around the genes with zero, low and moderate to high expression, reflecting the expected genomic activities around the genes. Overall, relative enrichment or depletion of labels is low for the genes that are not expressed compared to the expressed genes, while for genes with moderate to high expression suppressed labels are depleted and active labels are highly enriched. Transcribed label, for example, is enriched at moderate to low levels for genes with moderate to high and low expression, and it is mildly depleted for genes that are not expressed. On the contrary,FacultativeHet, associated with tissue-specific repression, is highly depleted among the expressed genes, and has moderate enrichment at genes that are not expressed.

**Figure 2:**
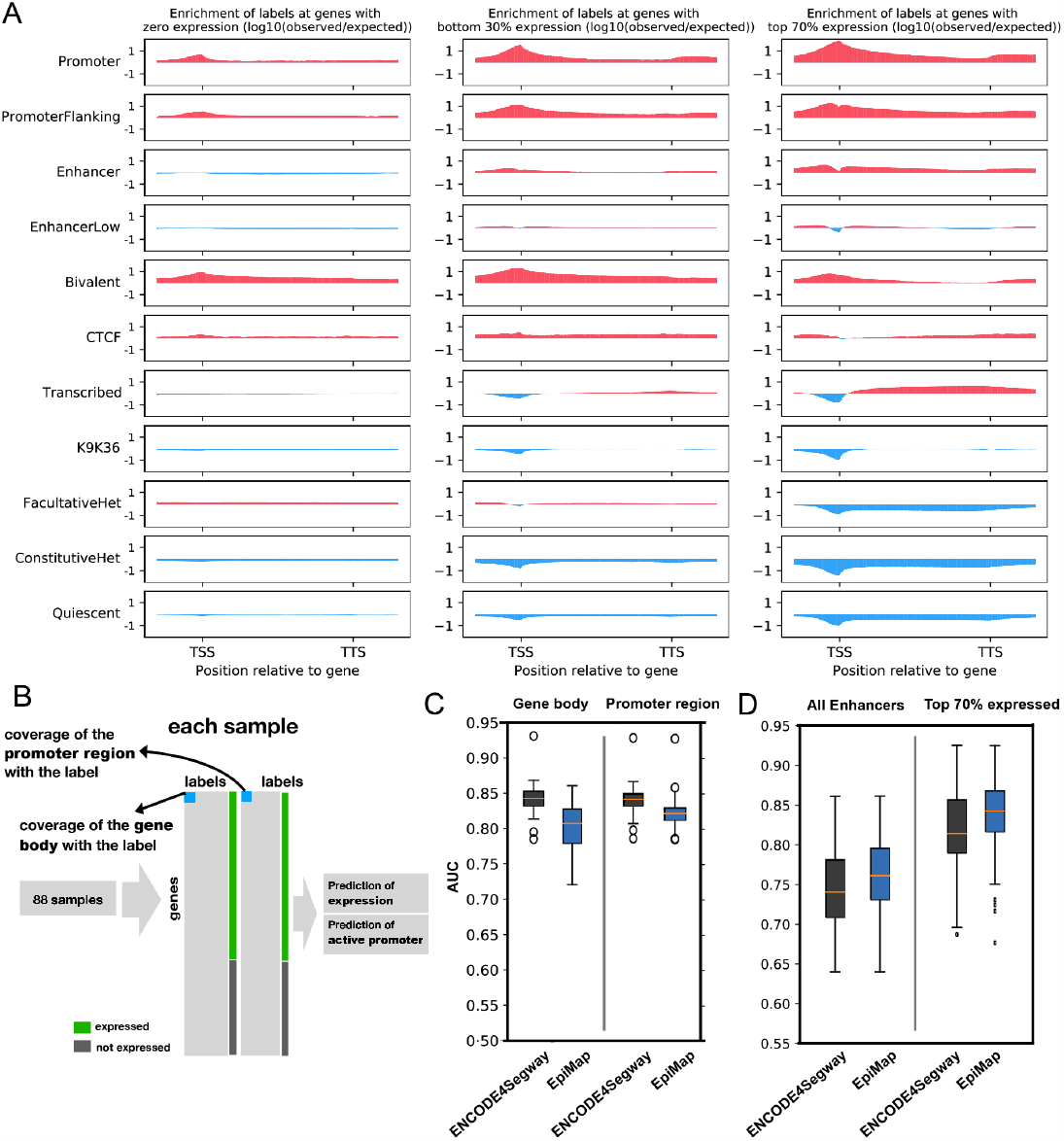
(A) Enrichment of the labels around genes, divided by expression level. Vertical axis indicates the degree to which the label occurs more at a given position than would be expected by chance if labels were distributed randomly (in the case of negative enrichment, less than expected by chance). (B) Pipeline for prediction of gene expression and active promoter regions based on the annotations (Methods). (C) Annotations predict gene expression and active promoter with high AUC. Vertical axis indicates the area under the receiver-operator curve (AUC) for predicting RNA-seq expression from labels at the gene body (left) and promoter (right) respectively (Methods). (D) Same as (C), but for predicting transcribed enhancers (eRNA production).

**Figure 3:**
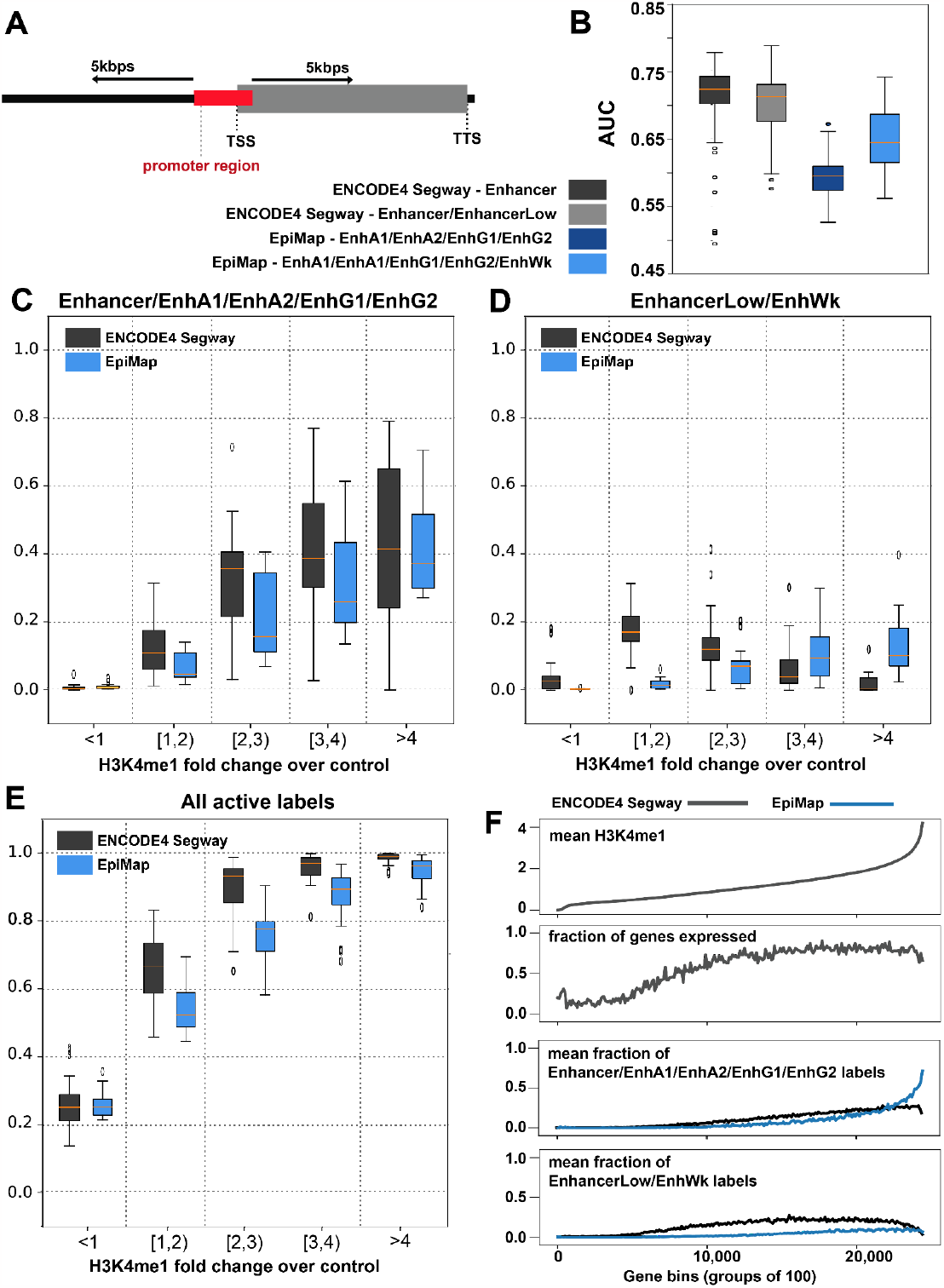
(A) Regions within 5kbps around genes TSS were selected for examination of Enhancer activity, excluding 2300bps (2000bp upstream, 300bp downstream of the TSS) promoter region (Methods). (B) Prediction of the gene expression based on the coverage of Enhancer labels in the 5kbps regions, for both EpiMap and ENCODE4 Segway annotations (Methods) . (C) Boxplots showing the coverage of Enhancer for ENCODE4 Segway annotations and EnhA1/EnhA2/EnhG1/EnhG2 for EpiMap annotations as a function of H3K4me1 signal. (D) Similar to C, but for EnhancerLow for ENCODE4 Segway and EnhWk for EpiMap annotations. (E) Similar to C/D, but including all active labels. (F) For one sample (ENCODE annotation accession ENCSR388IAJ), from top to bottom: first plot shows mean H3K4me1 values surrounding (5kb up/downstream, excluding 2300bp promoter region) each gene, for bins of 100 genes, sorted by this mean H3K4me1 value. Second plot shows the fraction of genes that are expressed (TPM>0). Third and forth plots show the mean fraction of coverage for labels Enhancer/EnhA1/EnhA2/EnhG1/EnhG2 and EnhancerLow/EnhWk respectively for the same region around each gene.

To quantitatively assess these annotations, we evaluated their efficacy towards understanding tissue-specific gene regulation by performing three analyses.

First, we evaluated how well Segway annotations of promoter activity capture gene transcription. Following previous work^7,12^, we used logistic regression to predict sample-specific gene expression based on either (1) the Segway labels at the gene’s promoter and (2) labels at its gene body, for a set of 88 samples with transcriptomic data available (Figure 2B, Methods).We found that the labels within each region are strongly predictive of expression (median AUC 0.84), indicating that the annotations accurately capture gene regulation. Our annotations are more predictive than those of EpiMap for both the gene body (median AUC 0.84 compared to 0.81 for Epimap), promoter regions (median AUC 0.84 versus 0.82). We also examined the association between Enhancer annotation labels and enhancer eRNA transcription measured by the FANTOM5 consortium ^18,19^, and although both Segway and EpiMap are predictive, their difference is not statistically significant (Figure 2D, see Methods).

Similarly, we found that Enhancer labels distal to a gene’s TSS are predictive of gene expression. We used the coverage of Enhancer labels within the 5kbp of the gene TSS and TTS as a predictor of gene expression, excluding 2000 bp upstream and 300 bp downstream around the TSS area for the promoter activity (Figure 3A, Methods). Results are similar for alternative choices of distal regions (see SFigure 7). Enhancer activity identified by ENCODE4 Segway is a predictor of gene expression with median AUCs of 0.72, 0.72, 0.69 for the three extended regions among the 88 samples (see Figure 3B). In comparison, EnhancerLow activity is not a strong predictor of gene expression (median AUCs of 0.59, 0.62, 0.6). EpiMap coverage of the combination of Enhancer labels (EnhA1, EnhA2, EnhG1, EnhG2, see SFigure 2 for mapping of labels) is a much weaker predictor of gene expression for most samples (median AUCs of 0.56, 0.59, 0.6, highest value 0.67). The combination of Enhancer and EnhancerLow state labels resulted in slightly lower prediction values for ENCODE4 Segway and higher prediction values for Epimap annotations.

We hypothesized that our higher accuracy at detecting transcription-associated activity derives from the use of genomic signals rather than binarized peak calls, leading to increased sensitivity for enhancer activity. To evaluate this hypothesis, we evaluated the sensitivity of each annotation set for enhancers as a function of H3K4me1 signal, a canonical mark of enhancers. Specifically, we examined the coverage of ENCODE4 Segway labels as a function of the intensity of H3K4me1 fold change values. As expected, we found that positions with higher H3K4me1 values are more likely to be annotated as Enhancer: 42% of positions H3K4me1 values >4 are labeled Enhancer (and have some kind of active label nearly 100% of the time), whereas only 0.4% of those with H3K4me1 values of 0-1 are labeled as Enhancer (Methods, Figure 3C, SFigure 08 for examples of individual samples). A similar pattern holds for EpiMap. However, whereas positions with moderate H3K4me1 signal (2-3 fold change) are rarely labeled as any type of enhancer by Epimap (median 0.16 EnhA1/EnhA2/EnhG1/EnhG2 and 0.07 EnhWk respectively), our annotations usually label such positions are as EnhancerLow (0.36 for Enhancer, 0.12 for EnhancerLow; Figure 3C). Finally, we observed that the coverage of the Enhancer label around the gene body and the mean intensity of the signal is a predictor of gene expression for individual samples (Figure 3D, SFigure 8). Our results show that ENCODE4 Segway Enhancer annotations are a good representative of moderate-to-high H3K4me1 fold change signal. We have also shown that the coverage of the Segway Enhancer label around the gene body is a predictor of gene expression.

### Annotations identify meaningful disease-cell type associations

We hypothesized that each variant-phenotype association identified by a genome-wide association study (GWAS) is driven by a functional genomic element which is active in a subset of cell types. Due to linkage disequilibrium, a GWAS can identify the phenotype-associated genomic regions, but not the exact SNP ^21,22^. Other works have demonstrated how chromatin state annotations can be utilized to identify both the functional elements driving the associations and the cell types in which these functional elements are selectively active ^7,20,23–30^. Here we use Segway annotations and conservation-associated activity scores (CAAS) ^7^ to identify cell type-specific functional genomic elements associated with GWAS traits.

We assigned the conservation-associated activity score (CAAS) to each chromatin state label to distinguish putatively functional types of activity (e.g. active regulatory elements or transcribed genes) from putatively nonfunctional types (e.g. inactive heterochromatin).Briefly, following previous work ^7^, we calculated the CAAS of each label using the phyloP conservation scores ^31^ of genomic positions which were annotated by that label, taking the seventy fifth percentile of absolute phyloP values at these label-associated positions as the label’s CAAS; higher CAAS indicates that label-associated genomic loci tended to have more extreme levels of conservation activity, which could in turn be interpreted as indicating a higher degree of biological functionality. Label CAAS was computed separately for each biosample because Segway produces sample-specific maps of regulatory activity from epigenetic signal tracks.

CAAS is a fully data-driven estimate of putative functionality; it is an alternative to manually choosing a subset of labels (e.g. Promoter and Enhancer) as putatively functional. A data-driven approach is important because different cell types may exhibit different functional activity; for example, functional regulatory elements may be poised in embryonic cell types and active in developed cell types. Notably, while the CAAS of labels at a given genomic position is correlated with evolutionary conservation, it represents an orthogonal measure of functionality to conservation, as it can identify human-specific functional elements. Note that we performed all analysis at the level of Segway state labels, not interpretation terms, to avoid potential bias introduced by the interpretation process.

We found that our functional and nonfunctional states are effectively distinguished by the CAAS value. CAAS was consistently higher for states receiving functional interpretation terms in the automated annotation pipeline (Figure 4A), with Promoter, Enhancer, Transcribed, and Bivalent states typically displaying the highest CAAS and Quiescent, ConstitutiveHet, FacultativeHet, and K9K36 states typically displaying the lowest CAAS. Furthermore, labels which received the same interpretation terms generally displayed similar but not identical CAAS, (SFigure 08), allowing downstream analysis to distinguish identically-interpreted labels which exhibit subtle differences in functionality (Methods).

**Figure 4:**
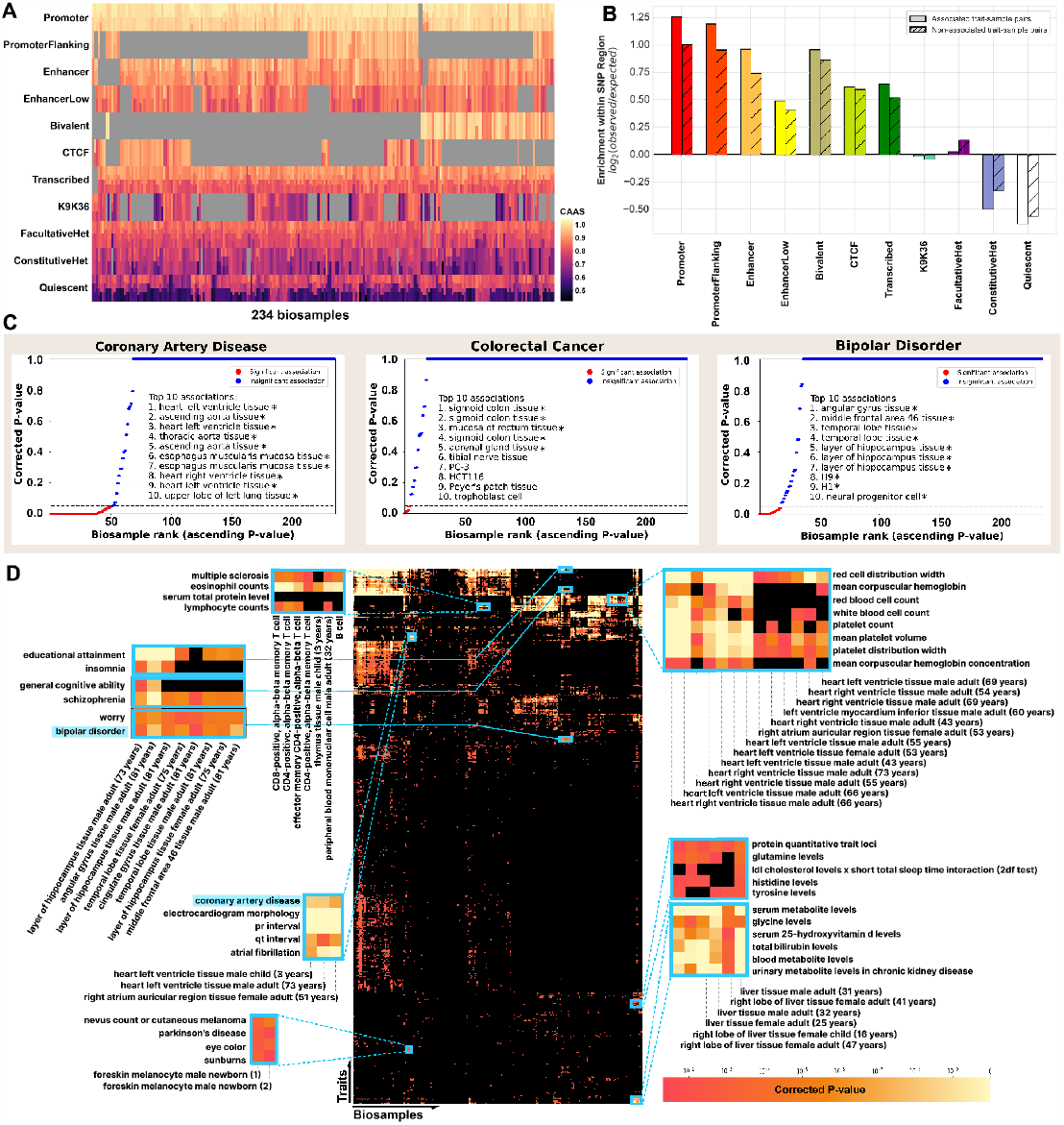
(A) Heatmap showing the CAAS values for the labels for 234 biosamples. Samples with multiple colors per row contain multiple labels that are assigned the same interpretation term. Gray cells indicate that the given sample does not have any labels with the corresponding interpretation term. (B) Enrichment of each interpretation term’s coverage within trait-associated SNP regions relative to the term’s coverage within the whole genome. Traits are grouped based on their association with each biosample; (trait, biosample) pairs which have a significant association (P < 0.05, “Associated trait-sample pairs”, solid bars), and pairs which do not have a significant association (“Non-associated trait-sample pairs”, hachured bars). (C) P-values for the association between three selected traits and the 234 biosamples. The 10 biosamples with the highest association (smallest P-values) are included in the plots. (*) indicates a significant association. (D) P-value matrix for associations between a subset of traits (rows) and the 234 biosamples (columns). The matrix is clustered along both axes. Non-black cells represent significant associations between a (trait, biosample) pair.

Given the effectiveness of our annotations at capturing biological functionality, we used the predicted sample-specific maps of regulatory activity to identify putative causal loci of GWAS associations ^32^ by examining the functional elements in the vicinity of their associated SNPs (Methods). We hypothesized that, if a given cell type plays a role in a trait, functional genomic elements involved in the trait are active in that cell type and thus GWAS SNPs are located near such elements. To evaluate this hypothesis, for each trait studied with a GWAS in the NHGRI-EBI GWAS Catalog, we first measured the significance of association between each trait-sample pair, then compared associations by ranking and clustering to assess their quality relative to known biology (Methods). We found that, as expected, GWAS SNPs are highly enriched nearby active regulatory elements, suggesting that such elements are good candidate causal drivers of disease association (Figure 4B). Furthermore, this enrichment is particularly strong in samples identified to be associated with the trait in question (Figure 4B, following paragraphs).

To identify samples involved in GWAS traits, we computed the mean label CAAS in the region surrounding trait SNPs for each sample, obtaining a metric which quantified the degree of functional activity in the vicinity of each SNP within that sample (Methods). We then used mean CAAS to rank the functional activity around every SNP across all samples, and used the Wilcoxon signed-rank test to assign a P-value to each trait-sample pair based on the ranks of that trait’s SNPs within the sample. A small P-value for a trait-sample pair indicated that trait SNPs exhibited more regulatory activity in the sample relative to the same trait in other samples and relative to different traits in the same sample (Methods). Relative to existing approaches for identifying cell type-disease associations ^29,33–39^, this approach has the advantage that each identified sample-trait association is supported by putative driver elements with measured activity across 234 samples.

The test for differential trait-sample associations resulted in 16,127 significant (Bonferroni-corrected p<0.05) trait-sample associations after correcting for multiple testing (Methods). SNP regions from significant associations were more enriched in functional elements than SNP regions from insignificant associations, as expected from the way the test was designed (Figure 4B). Conversely, in samples not associated with the trait in question, SNP regions are more likely to receive FacultativeHet, suggesting that the activity of such loci is often specific to associated samples and that they are repressed in non-associated samples (Figure 4B).

Statistical testing for differential association revealed associations between traits and cell types that align with known biology. For example, the five biosamples most associated with the coronary artery disease trait were all heart tissues; three of the five significant associations with the colorectal cancer trait were colon tissues (the remaining two being rectum cells and adrenal gland cells); and the top seven significant associations with the bipolar disorder trait were brain tissues and cell types (Figure 4C). Additionally, hierarchical clustering based on P-values resulted in co-clustering of similar cell types from different donors and in co-clustering of similar traits, reinforcing the quality of our annotations (Figure 4D). Interestingly, the Parkinson’s disease trait clustered with traits for eye color, sunburn, and cutaneous melanoma, and the entire cluster of traits exhibited significant association with two melanocyte samples; these associations again align with known biology, as multiple studies have shown a link between Parkinson’s disease and melanoma ^40^. Traits with a huge number of associated SNPs (many hundreds or thousands) have significant p-values for many samples; these traits tend to be those, such as “educational attainment”, with likely very complicated biology and potential for bias in data collection (Figure 4D, top right corner; see STable 04 for complete list of significant associations).

## Discussion

Here we present the ENCODE4 Segway Encyclopedia, a collection of sample-specific chromatin state annotations produced using the Segway pipeline. We showed here that these annotations comprehensively summarize epigenomic data from each sample and accurately capture many known genomic phenomena including gene regulation and regulatory elements. We have distributed these annotations through the ENCODE portal, which makes them easy to organize, view and download. As SAGA chromatin state annotations are a simple and easy-to-use summary of a large collection of data, we expect that these annotations will provide an easy entry point for researchers looking to make use of epigenomic information.

The ENCODE4 Segway Encyclopedia has a number of advantages over alternative annotations of genomic elements. Unlike annotation strategies that consider only a single mark, this encyclopedia is integrative, and thus the annotated elements are informed by all epigenomic data sets measured in the target sample. Relative to existing large-scale SAGA annotations, we showed that the ENCODE4 Segway Encyclopedia has increased sensitivity, likely due to its use of genomic signals.

We demonstrated here that the ENCODE4 Segway Encyclopedia enables researchers to identify causal elements driving disease- and phenotype-associated genetic variation. We found that observed disease-associated genetic variants can usually be explained by a putatively functional genomic element within a typical linkage disequilibrium window of that variant. Furthermore, we showed that doing so can elucidate the cell types involved in disease.

An important caveat of this analysis is that all of ENCODE’s epigenetic data sets are derived from bulk samples and thus the resulting annotations do not necessarily represent the activity of any one cell. This is particularly important to consider when considering the difference between low- and high-signal loci. It is an open question whether low signal at a locus represents a homogeneous set of cells with weak activity or a heterogeneous set of cells, some with strong activity and some with no activity. As single-cell data becomes available, it will increasingly become possible to untangle this open question, and perhaps even to produce chromatin state annotations of single cells or homogeneous sets of single cells.

## Data Availability

Each of the 234 ENCODE4 Segway annotations are available on the ENCODE portal in a bed9+ file format. These genome-wide annotation files include the coordinates of genomic regions, their chromatin state label, the RGB color used for that state label in the genome browser and the label initially generated by Segway. The metadata for each sample, as well as the list of the track files which were used to generate the annotations and visualization on the ENCODE genome browsers are also available in the ENCODE portal (https://www.encodeproject.org/report/?type=Annotation&lab.title=Maxwell+Libbrecht%2C+SFU&field=accession&field=files&field=files.status&limit=200&status=released).

The ENCODE portal includes a unique accession for each annotation along with all relevant metadata. This metadata includes all datasets from which the annotation is derived, each with an accession and link to publicly-available raw data. It also includes the identity and version number of every tool used in the pipeline from input to output.

On the ENCODE portal, we have also included a set of sample-specific plots demonstrating the properties and statistics of each annotation. For each sample, four plots demonstrate the mean signal value of the input tracks, the classifier probabilities from the interpretation process, the emission probabilities and the genome coverage for each of the labels. A fifth plot demonstrates enrichment of the labels around the gene body. For samples with transcriptomic data available, this plot has sections for genes with zero expression, bottom 30% expression and top 70% expression (similar to Figure 2).

The annotation pipeline code is available at:

https://github.com/ENCODE-DCC/segway-pipeline/

Code for result sections 1 and 2 is available at: https://github.com/marjanfarahbod/SegwayClustering

Code for result section 3 is available at: https://github.com/ardiab/encode4_segway_catalog_gwas

## Methods

### Datasets

We annotated all ENCODE4 cell types with sufficient epigenomic data. We selected a panel of data sets that were available in most cell types. Specifically, we used six ChIP-seq measurements of histone modification H3K4me3, H3K4me1, H3K27ac, H3K36me3, H3K27me3, H3K9me3; DNase-seq or ATAC-seq measurements of open chromatin; and ChIP-seq measurements of CTCF binding.

We processed each sequencing data set into a real-valued tracks using the ENCODE uniform pipelines ^41^. Briefly, reads were mapped to the human reference genome; reads were extended according to inferred fragment length. For ChIP-seq data, we applied a fold enrichment normalization by dividing the observed signal by input control signal.The output of this processing is a track over the genome that assigns a real-valued signal strength to each genomic position.

We chose to annotate all samples with at least the six histone marks listed above; all annotated samples have these marks, but some are missing DNase/ATAC-seq or CTCF. Only a few have POLR2A and EP300. When multiple data sets for the same (cell type, assay) pair were available, we chose the more recently processed data.

For input to Segway, we binned signal data sets at 100 base pair resolution by taking the average for each 100 bp bin. We excluded unmappable positions and the ENCODE exclusion list; these are considered to be unobserved by Segway (see below). We applied the variance-stabilizing inverse hyperbolic sine transform 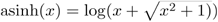 to all signal data sets.

For transcriptomics-based evaluation (see below), we used RNA-seq data for all annotated cell types where this data was available. In total, 88 of samples (38%) had the transcriptomic data available. For each of these samples, the “Total RNA-seq - Default - gene quantifications’’ was downloaded from ENCODE portal, matching the tissue and the donorID.

We acquired gene coordinates from https://www.encodeproject.org/files/gencode.v29.primary_assembly.annotation_UCSC_names for the 26,017 genes.

For enhancer eRNA evaluation (see below), we acquired Cap Analysis of Gene Expression (CAGE) data from FANTOM5 ^18,19^ (https://fantom.gsc.riken.jp/5/datafiles/latest/extra/Enhancers/). These data sets measure eRNA transcription for 65,423 potential enhancer regions in 1,828 cell types. We were able to match 175 of our samples to the same tissue type in the FANTOM5 data. Supplementary Table ST02 contains the list of matched cell types.

EpiMap annotations ^20^ were obtained from the portal for all the Segway annotations from the same cell type/tissue. When available, we used the EpiMap annotations from the matching donors. Supplementary table ST03 contains the list of matching EpiMap and Segway accessions.

### Annotation interpretation

For the interpretation process, Random Forest classifier was used according to Libbrecht et al. 2019 ^7^. We trained the classifier using the available training data from the same reference, (count of 210 states) and a set of 90 states from the new set of runs. We selected the new interpretation term set based on these two sets of data.

### Evaluation based on prediction of transcription

For the 88 samples with the transcriptomic data available, we used coverage of the annotation states on the gene body and promoter regions to train and test the logistic regression to determine the expression of the genes (STable 01 includes the list of transcriptomic data). We obtained gene coordinates from https://www.encodeproject.org/files/gencode.v29.primary_assembly.annotation_UCSC_names/ and considered 2000 upstream and 300 downstream of the TSS as the promoter region. For each sample, we trained the classifier on 80% of the genes and tested on the remaining 20%.

For prediction of gene expression based on the coverage of Enhancer/EnhA1/EnhA2/EnhG1/EnhG2 labels, we used the fraction of coverage of these labels at regions within 2kbps, 5kbps and 10kbps around the gene TSS (excluding the 2300bps promoter region) as the predictor. (see STable 03 list of EpiMap annotations obtained from the ENCODE portal)

### Evaluation based on enhancer RNA transcription

We first converted the enhancer coordinates to match the genome assembly GRCh38/hg38 using the genome assembly converter available in http://genome.ucsc.edu ^42^. Similar to the transcription analysis, for the 175 samples with a matching tissue in FANTOM5 eRNA transcription ^18,19^, for each sample we trained the logistic regression on the 80% of the potential enhancer regions and tested on the remaining 20%. The mapping of our samples to FANTOM5 samples is available in STable 02.

### GWAS SNP analysis

#### Processing SNPs from the EBI GWAS catalog

We obtained the locations of 209,555 unique trait-associated single nucleotide polymorphisms (SNPs) identified by 5,197 genome-wide association studies (GWAS) from the NHGRI-EBI catalog of human genome-wide association studies ^32^. The SNPs were associated with 15,143 different traits, and were obtained after removing null entries from the catalog. We then applied four preprocessing steps to the GWAS SNPs.

First, we excluded all trait-associated SNPs which fall within the human MHC genomic region. Second, we replaced each GWAS SNP with an associated “SNP region” to include neighboring genomic positions which may be in LD with the GWAS-identified polymorphism and which may underlie the disrupted biological process causing the observed phenotype ^43^. SNP regions were defined as 20,001 bp genomic windows centered on the associated SNP, and were clipped on the appropriate side when a SNP was located less than 10,001 bp from one end of the chromosome. Third, we filtered the SNP regions associated with each GWAS trait to prevent double-counting. Specifically, we sorted the SNP regions associated with each trait in order of ascending SNP P-value and added regions greedily to the trait; a SNP region created around a SNP with a larger P-value was only added to the trait if it had an intersection of less than 50% with each previously-added trait SNP region that was created around a lower P-value SNP. We note that this processing step was done at the trait level, meaning that a given SNP which is associated with multiple traits could be filtered out for only a subset of the traits. In the fourth and final SNP processing step, we removed traits with fewer than 30 filtered SNP regions.

After the preprocessing steps, 144,071 unique SNP regions and 1,274 traits remained.

#### Measuring enrichment of functional elements in SNP regions

To measure the level of functional activity within SNP regions, we intersected the 144,071 filtered SNP regions with Segway annotations for each of the 234 available biosamples, using the bedtools package ^44^. The intersections yielded chromatin state distributions for every (biosample, SNP region) pair, which we then used to calculate a biosample-specific metric (mean CAAS) that encodes the functional activity of each SNP region in each biosample.

#### Calculating label-wise conservation-associated activity score (CAAS)

We calculated the conservation-associated activity score of each Segway label in each biosample following Libbrecht et al. 2019 ^7^. We used phyloP scores ^31^ derived from the genomes of 30 mammals, 27 of which are primates (download link https://hgdownload.cse.ucsc.edu/goldenpath/hg38/phyloP30way/hg38.30way.phyloP/)

Let *L*_*jk*_ represent the set of all genomic bins *i* assigned label *j* by Segway in biosample *k*. Then, the CAAS of label *j* in biosample *k* is defined as:

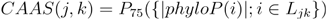

where *P*_75_ denotes the 75th percentile and *phyloP(i)* denotes the phyloP score at position *i*. As previously observed, we found that the 75^*th*^ percentile achieved good separation in phyloP between labels (SFigure 9).

#### Calculating mean CAAS in a SNP region

We measured the enrichment of regulatory activity within the SNP regions using their mean CAAS. Given a SNP region *R=* { *i*_1_,…,*i*_20,001_ } and a chromatin state annotation *Ann*(*i,k*):{*i*∈[1,2,…]}→{*j*∈[1,…,*J*]}which assigns a label *j* to each position *i* in biosample *k*, the mean CAAS for the region in a given biosample *k* was obtained by taking the average of the label CAAS within the biosample, weighted by the proportion of the SNP region covered by each label in that biosample (SFigure 9):

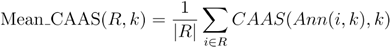

where |*R*|=20,001 is the number of bases in the SNP region.

Intuitively, a high mean CAAS within a SNP region indicates that the region received Segway chromatin state annotations which were used to annotate highly-conserved positions in the genome; consequently, using conservation as a proxy for functional importance, such regions can be thought of as having a high degree of functional activity.

#### Testing for differential biosample-trait association

We used the mean CAAS values for SNP regions to test for differential association between (biosample, trait) pairs. For each of the 144,071 SNP regions *S* ={*s*_1,_…,*s*_144,071_ }defined in the SNP preprocessing step, we ranked the 234 biosamples *B* = {*b*_1,_…,*b*_234_} by the CAAS of the SNP region, producing ranks between 1and 234, where a rank of 1 signifies that the biosample had the lowest CAAS for the SNP region, and a rank of 234 signifies that it had the highest CAAS. The overall ranking process produced a ranking matrix *R* ∈ ℕ^144,071×234^, where row *i* contains the CAAS ranks of all biosamples *b*_*j*_∈*B* for SNP region *s*_*i*_, column *j* contains the CAAS ranks of biosample *b*_*j*_ for all SNP regions *s*_*i*_∈*S*, and entry (*i,j*) specifies the CAAS ranking of biosample *b*_*j*_ for SNP region *s*_*i*_.

We then calculated a “null” rank for each biosample by taking the median rank across all SNP regions for the biosample; the null rank for biosample *b*_*j*_ is calculated as

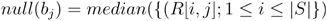

To test for the degree of association between a biosample *b*_*j*_ and a trait *T*_*k*_ ={*s*_*m*,_…,*s*_*n*_} ⊂ *S* that is associated with a subset of the SNP regions, we used the Wilcoxon signed-rank test to test whether the median rank within the specific trait’s rank distribution was greater than the null rank of the biosample. Intuitively, the test asks the question

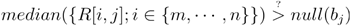

Biosample-specific null ranks were used as a normalization method to control for biosamples which exhibit high overall regulatory activity, so that the test can better capture differential association in biosamples which exhibit lower overall regulatory activity that is more specific in nature.

The test produced a P-value matrix 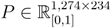, where an entry (*k,j*) specifies the P-value for the association between trait *T*_*k =* {_ *s*_*m*_,…*s*_*n*_ }⊂ *S*, and biosample *b*_*j*_. Lower P-values can be interpreted as specifying higher-than-expected ranks for the trait’s activity within the biosample, and therefore as indicating a more significant association between the (trait, biosample) pair. To control for multiple testing, we applied Bonferroni correction by multiplying all P-values by the number of biosamples. STable 04 includes the list of significant (biosample, trait) associations with the corresponding p-values.

#### Hierarchical clustering

To investigate whether the obtained (trait, biosample) associations captured biological differences, we clustered the P-value matrix produced by the test for differential biosample-trait association. If the test detects real biological patterns, traits which share underlying functional mechanisms are expected to co-cluster, whereas traits with differing mechanisms are expected to appear in distinct clusters (indicating that the test successfully assigned broadly similar P-values to similar traits and different P-values to dissimilar traits). Analogously, biosamples consisting of similar cell types are expected to co-cluster.

We applied Euclidean hierarchical clustering to the P-value matrix produced by the test for differential biosample-trait association. Our approach applied clustering along both axes of the matrix (traits and biosamples) based on the computed P-values.

