## supplemental Material for "Integrative chromatin state annotation of 234 human ENCODE4 cell types using Segway reveals disease drivers"

### Supplementary material
### Chromatin state annotation of cis-regulatory elements in hundreds of human cell types using Segway

#### Review of existing SAGA catalogs

Here, we briefly review the existing SAGA methods and describe their relationship to the ENCODE4 Segway catalog. (We recently reviewed these methods in more detail in [5].)

The first SAGA methods were developed in the 2000s and include HMMSeg, ChromHMM and Segway. Since then, a large number of SAGA approaches have been described (reviewed in [5]). Due to the challenges inherent in running at large scales, of those methods, only three—ChromHMM, Segway and IDEAS—have been applied at scale (100+ samples), and until now only ChromHMM has been applied at a large scale (200+ samples). There are two main differences that distinguish these three methods. First, ChromHMM uses binarized data as input to the SAGA method, while Segway and IDEAS use continuous signals. Second, the methods differ in their type of information sharing across samples (defined below): the Segway pipeline uses no sharing across samples, while the ChromHMM and IDEAS pipelines use both horizontal and vertical sharing.

SAGA methods for annotating multiple cell types can use two types of information sharing: horizontal and vertical sharing (reviewed in greater detail in [5]). Horizontal sharing emphasizes similarities for learning labels. The most commonly-used type of horizontal sharing is concatenated annotation, in which a single model is applied across all cell types. Horizontal sharing is widely used but has two downsides: (1) it requires that all samples have the same set of available data and (2) it can be vulnerable to sample-specific batch effects.

Vertical sharing emphasizes similarity for predicting the state of a given genomic position. Vertical sharing methods include TreeHMM, IDEAS, Segway+GBR and SPIN [9, 1, 8, 10, 4, 6]. Applying SAGA to imputed data (using an imputation method such as ChromImpute, PREDICTD, Avocado or others [7, 2, 3]) is another type of vertical sharing. Vertical sharing has the downside that the state in a given cell type can be influenced by activity in other cell types; in particular, these methods tend to mask differences between cell types [5].

The ENCODE4 Segway catalog uses neither type of sharing and thus each annotation represents an unbiased summary of the data in each cell type. That is, we learn a separate model for each cell type (no horizontal sharing) without imputation or otherwise referencing the data from other cell types (no vertical sharing). The only information sharing occurs at the interpretation stage, in which we use a single shared vocabulary to describe all states (a type of horizontal sharing).

#### Segway parameter setting

Segway train command:

```
segway train --max-train-rounds=25 --num-instances=10
       --track-weight=0.01 --segtransition-weight-scale=1
       --ruler-scale=100 --prior-strength=1 --resolution=100
       --minibatch-fraction=0.01 --num-labels=[number of labels - see below]
```

Number of labels is determined as $10 + 2\sqrt{M}$ where $M$ is the count of input datasets for the sample.

#### Process for identifying samples with low quality annotations

We investigated states with low prediction probability from the interpretation process and found that some samples have generally lower probability distributions for multiple states (see SFigure 04). Upon further investigation, we found that these samples tend to have high coverage of the Quiescent label from the

ChromHMM annotations (see SFigure 04). Determining a threshold for various metrics of such quality examination is challenging. Nonetheless, we decided to report these samples as well as keep a track of them in our downstream analyses. The quality of annotations can potentially explain some of the anomalies observed in their future application.

#### Supplementary figures

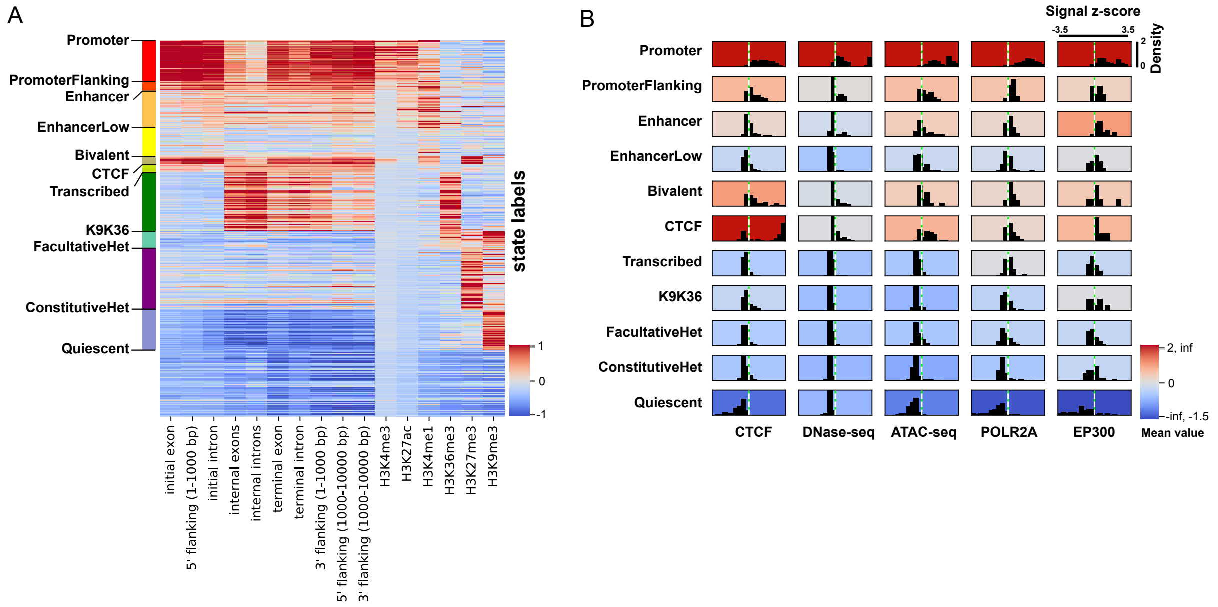


SFigure 1: (A) feature values for the state labels, grouped based on their assigned interpretation terms. (B) Distribution of the the five track values (not included in the feature set) for state labels from the samples which have these tracks.

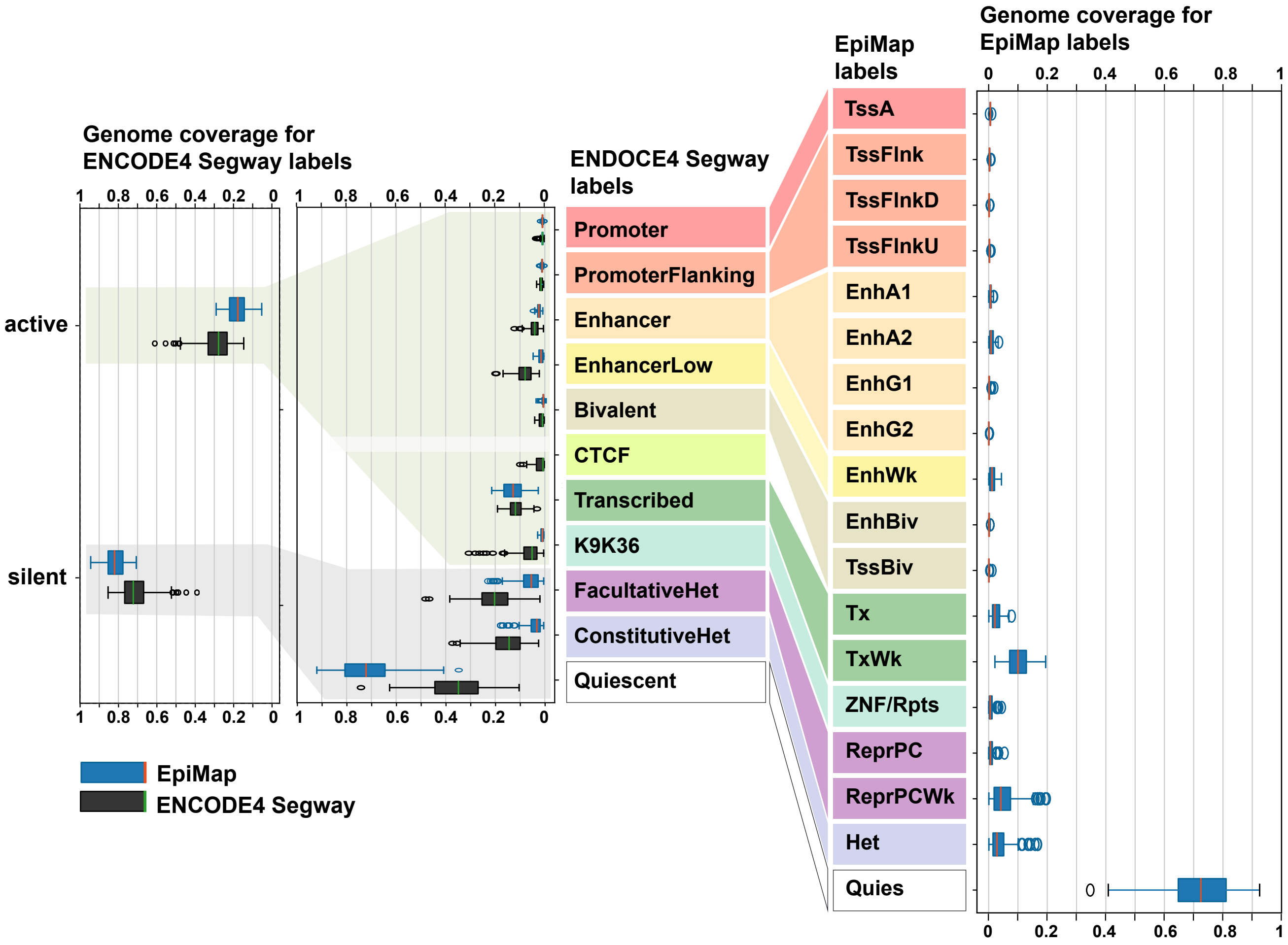


SFigure 2: Genome coverage of the labels by interpretation terms. Left: Genome coverage of the labels. EpiMap labels mapped to the ENCODE4 Segway labels. Right: genome coverage for EpiMap labels.

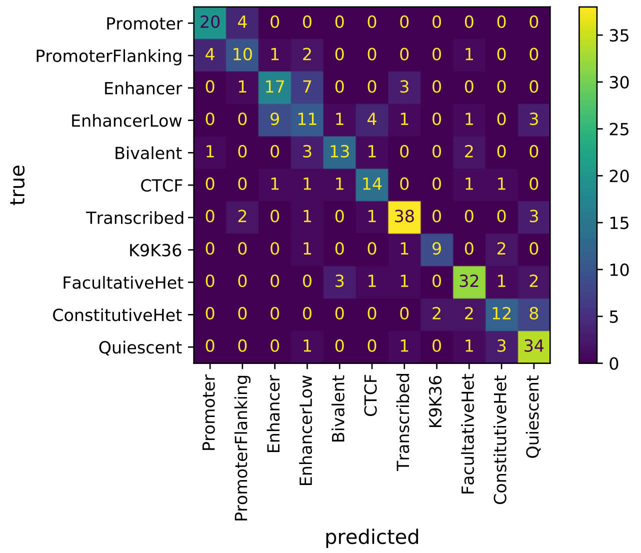


SFigure 3: Confusion matrix from the random forest 300 training samples.

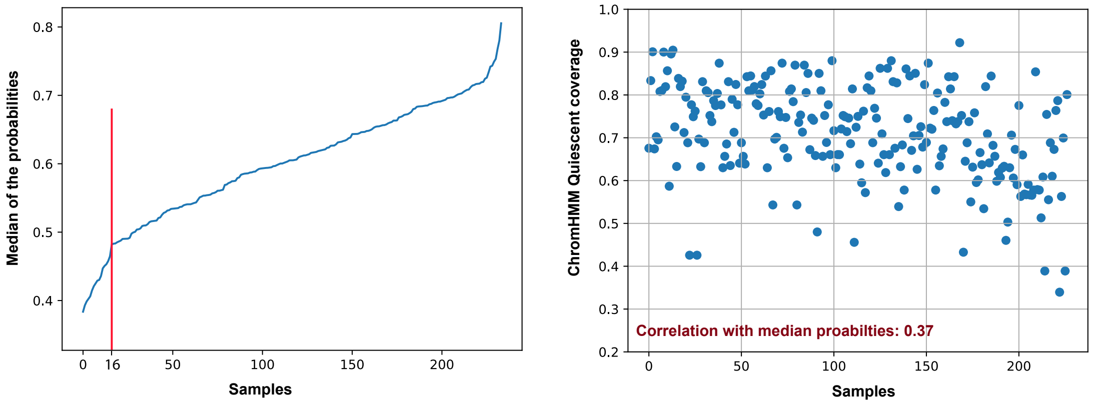


SFigure 4: Left: median probabilities from the interpretation process, sorted. Right: coverage of the Quiescent label by EpiMap annotations.

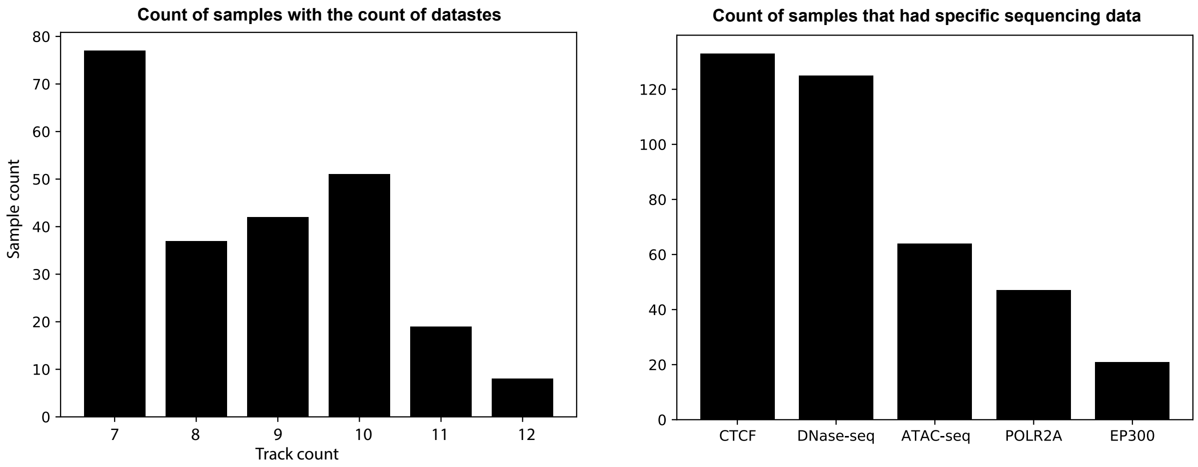


SFigure 5: Left: count of samples with different count of sequencing data. Right: count of samples with different sequencing data.

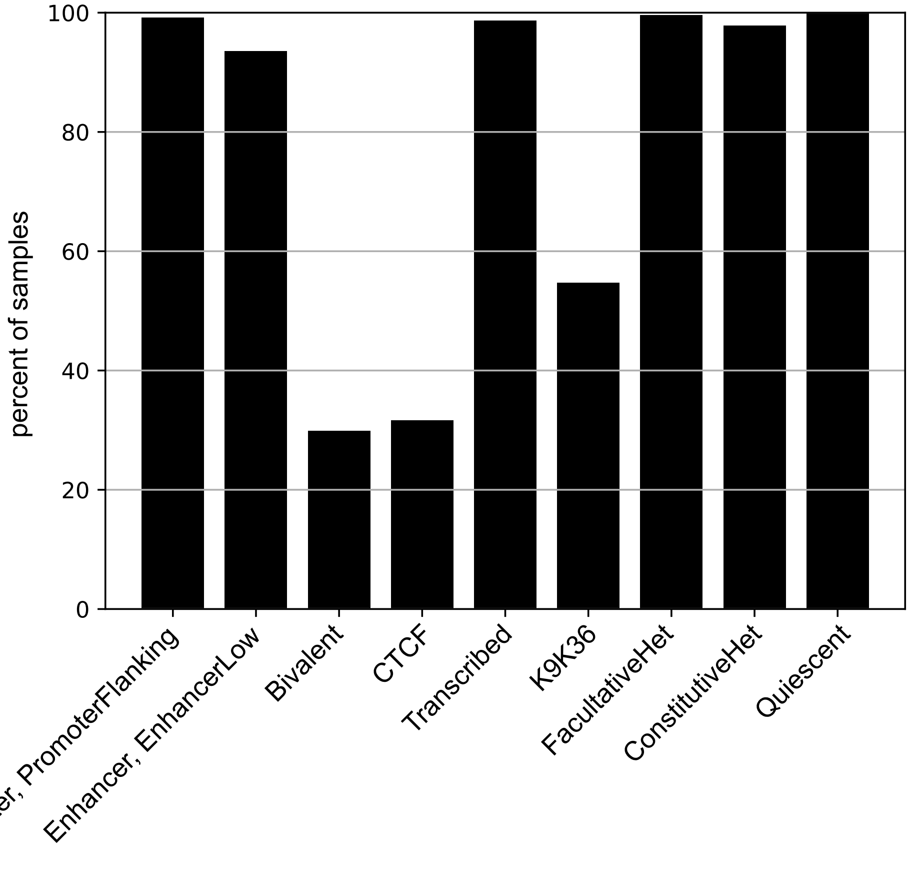


SFigure 6: Percent of samples with existing annotation states. Only 2 and 3 samples are missing Promoter and Transcribed labels, out of with 3 were identified as low quality samples based on the interpretation probabilities.

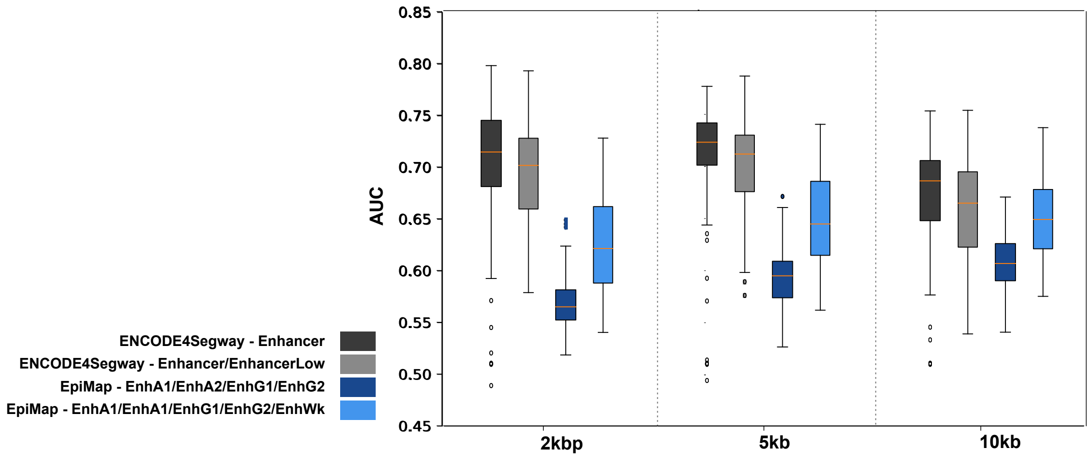


SFigure 7: Prediction of gene expression based on label coverage in regions within 2kbps, 5kbps and 10kbps around the TSS.

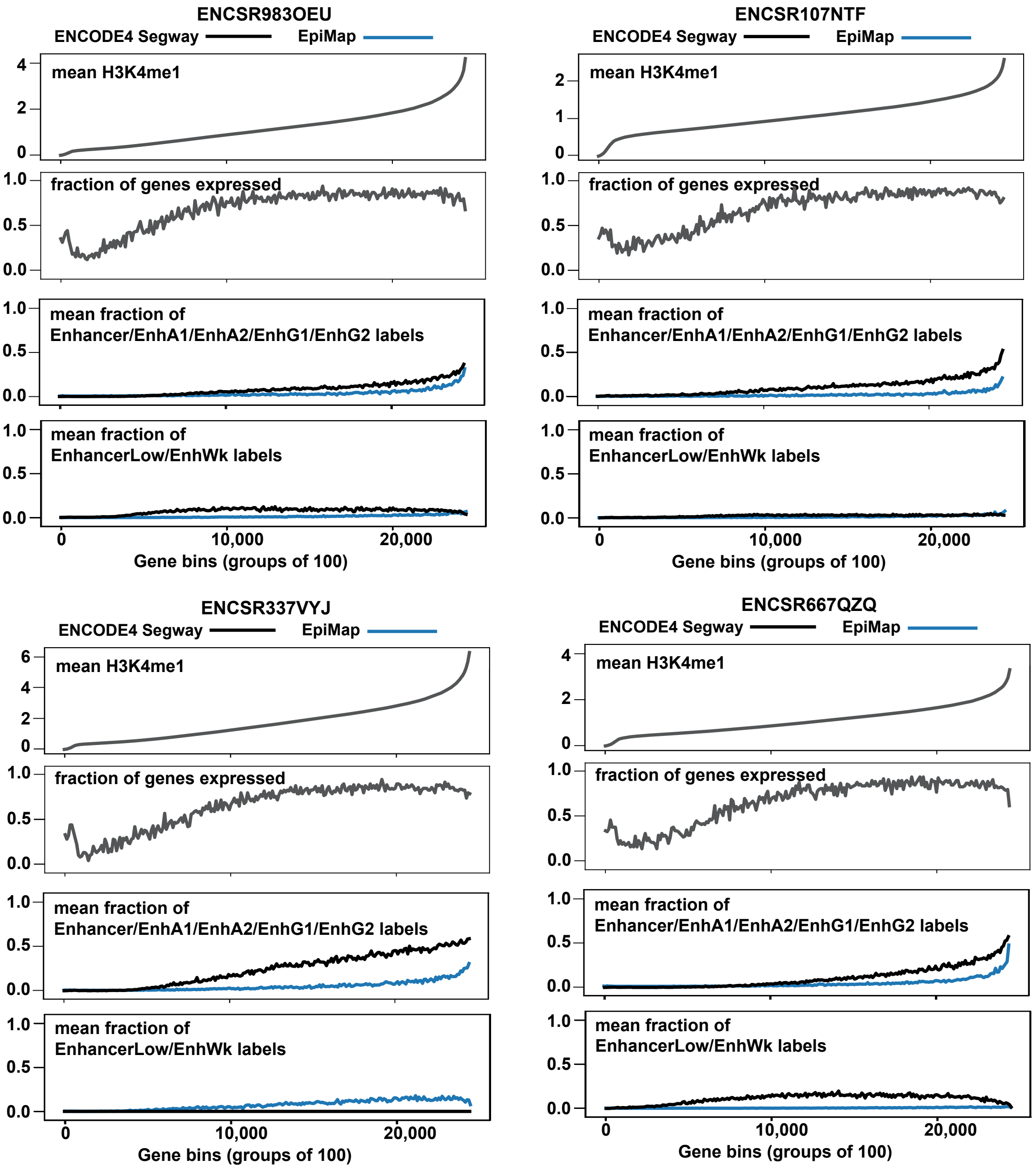


SFigure 8: More examples for Figure 3F. For each sample, from top to bottom: first plot shows mean H3K4me1 values surrounding (5kb up/downstream, excluding 2300bp promoter region) each gene, for bins of 100 genes, sorted by this mean H3K4me1 value. Second plot shows the fraction of genes that are expressed (TPM¿0). Third and forth plots show the mean fraction of coverage for labels Enhancer/EnhA1/EnhA2/EnhG1/EnhG2 and EnhancerLow/EnhWk respectively for the same region around each gene.

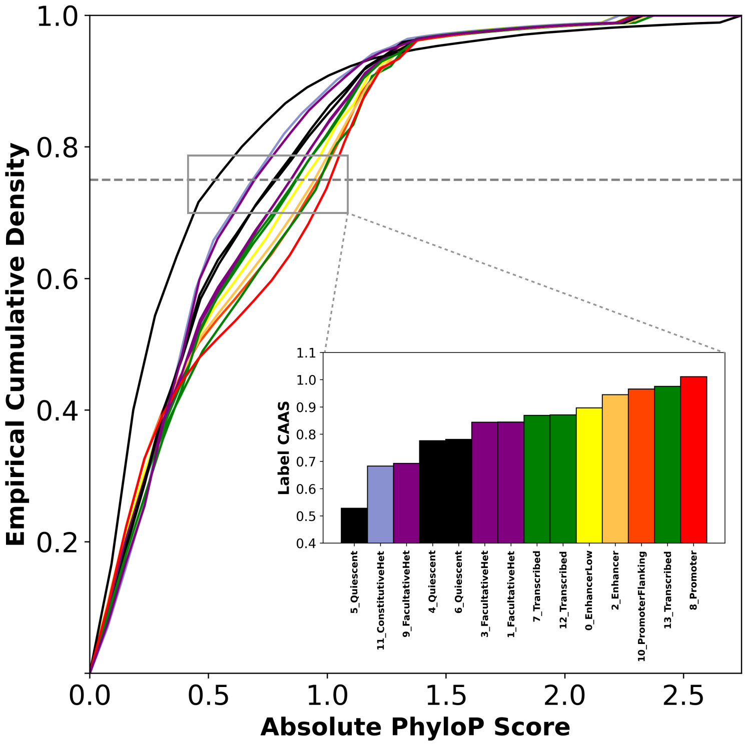


SFigure 9: Empirical cumulative densities of absolute PhyloP scores in label-associated genomic regions. The bar plot specifies the conservation-associated activity score (CAAS) associated with each label, which is obtained by taking the 75th percentile of the absolute PhyloP values at positions receiving the label as an annotation
